# Characterizing the frequency of non-allelic homologous recombination (NAHR) on the human X chromosome (Xq28) using digital inverted PCR

**DOI:** 10.1101/2020.04.20.051185

**Authors:** Lou Ann Bierwert, Samantha Bryce, Robert B. Merritt

## Abstract

Little is currently known about the rates at which non-allelic homologous recombination (NAHR) occurs. However, most current research suggests that NAHR is rare. Previous work by Small, et al (1998), examined an inversion polymorphism on the long arm of the X-chromosome, involving two genes (*FLNA* and *EMD*), and determined the frequency of the two gene arrangements in a group of European individuals. Here we quantify the rate at which the causal NAHR, in inverted repeats flanking the *FLNA* and *EMD* genes, occurs in meiosis using digital PCR of sperm samples, with male cheek cells as controls. NAHR was documented in all samples, including the cheek cell samples at a mean recombination rate of 1.8%, indicating that NAHR occurs much more frequently than initially believed, and appears to be occurring in mitosis. The increase in NAHR frequency in spermatogenesis is not significant leaving in question NAHR occurrence in meiosis. This study reveals a more accurate way to quantitate NAHR, serving as an important first step in better understanding various NAHR-associated diseases.

**Author Summary:** We sought to more accurately quantitate and characterize NAHR at a site at the end of the long arm of the X chromosome that contains a set of inverted repeats flanking two genes, *filamin* and *emerin*. We determined that NAHR is happening far more frequently than previously thought, and in this case unequally, depending on the direction of the inversion. We speculate on the possibility of local adaptation playing a role in this. These high-resolution results were obtained by modifying a previously published assay which can be easily adapted to other inversions. This could be especially helpful in studying those NAHR inversions related to disease.

## Introduction

Non-allelic homologous recombination (NAHR) is one of the main mechanisms for genomic rearrangements and is aided by low copy repeats (LCRs) (Gu, et al., 2008). The focus of this study is human NAHR frequency involving two 11.3kb long inverted repeats (LIRs) in the q28 region of the X chromosome responsible for a recurrent inversion with the identifier HsInv0389. The repeats associated with this inversion are greater than 99% identical and flank two genes, *emerin* (*EMD*) and *filamin* (*FLNA*) (Small, et al., 1997). The NAHR-driven inversion in this region results in two different gene arrangements (Fig 1); the plus orientation reads (centromere-*FLNA-EMD*-telomere), while the minus orientation reads (centromere-*EMD-FLNA-*telomere). Interestingly, LIRs have been found to flank *FLNA* and *EMD* in all eutherian mammals studied, with a single arrangement found in each species (probably a result of sample size) but both orientations being found with equal prevalence across species, indicating these LIRs act as hotspots for inversion breakpoints (Caceres, et al., 2007). In addition, in a sample of women of European ancestry, 33% were found to be heterozygous for the HsInv0389 inversion with no significant deviation from Hardy Weinberg equilibrium zygotic frequencies (Small, et al., 1998). Since both orientations are common in many human populations, it is possible to isolate single copies of both arrangements by using male subjects, making this particular inversion an ideal choice for better understanding NAHR.

**Fig 1.**
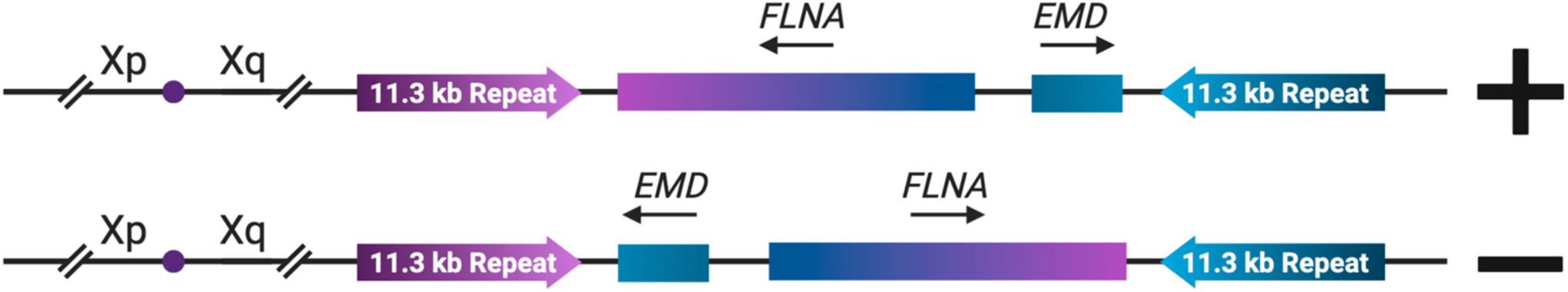
A graphical depiction of the two orientations of the HsInv0389 inversion. Located in the q28 region at the end of the long arm of the human X chromosome lies this inversion. The identical long inverted 11.3kb repeats drive this inversion, reversing the order of the Filamin and Emerin genes in this region. Adapted from Kirby et al., (2016).

While this *FLNA*-*EMD* inversion does not lead to any observable phenotypic effect, it is proposed that a rare unequal crossover event between inverted alleles results in the deletion of the *emerin* gene, causing Emery-Dreifuss Muscular Dystrophy (Kirby., et al. 2016). In addition, NAHR is associated with a vast array of deleterious genetic disorders including Smith-Magenis syndrome, Potocki-Lupski syndrome, pituitary dwarfism, and glabozoospermia (Carvalho and Lupski, 2016; Ellanti et al., 2012; Mefford, 2010). This prevalence of NAHR-related genomic disorders underscores the importance of improving our understanding of NAHR frequency, especially since it has also been the proposed major source of DNA damage associated with a range of cancers (Belancio, Roy-Engel, and Deininger, 2010; Gu et al., 2008). For example, early work identified recombination hotspots associated with the *BRCA1* and *BRCA2* genes, implicating these hot spots in breast cancer (Mazoyer, 2005). While it was unclear what recombination mechanism was being observed; Belancio, Roy-Engel and Deninger (2010) later realized that Mazoyer had actually tied NAHR to breast cancer. They also identified disruptions in genomic stability previously reported to be associated with repetitive elements and cancers such as colon cancer and retinoblastoma (den Hollander et al., 1999; Miki et al., 1992), demonstrating how NAHR and cancer biology are interwoven.

Given this ever-growing list of possible relationships between NAHR and human health, the need to fully understand the frequency and possible driving forces of NAHR is evident. Thus, this study sought to better quantify the NAHR frequency responsible for the HsInv0389 inversion with a modifiable assay that could be applied to other inversions of interest. Recent estimates of NAHR frequency are highly variable with one study finding a mean NAHR rate of 3.5×10^−5^ (MacArthur et al., 2014). However, due to the large size of most inverted repeats, in combination with popular genotyping techniques, and bias in favor of a reference genome orientation, identifying NAHR is likely often underestimated in current databases (Giner-Delgado et al., 2019). Using the iPCR method (Kirby et al., 2016), in combination with digital PCR allowed us to both more accurately detect NAHR occurrence in this study and refine methodologies that extend beyond the HsInv0389 inversion.

We initially hypothesized that this inversion only had an opportunity to occur during meiosis and more commonly in spermatogenesis where the X chromosome lacks a homologous chromosome with which it can recombine. However, Flores et al. (2007), demonstrated that inversions also occur in blood cells, indicating that somatic NAHR could produce mosaics of genomic structures within individuals. In this study, we report on the frequency of the HsInv0389 inversion during meiosis at this Xq28 site using digital iPCR on sperm samples. Through this process, we determined that NAHR is occurring at much higher rates than previously reported. In addition, we inadvertently confirmed that NAHR is also occurring during mitosis, as our “control” cheek cell samples also exhibited NAHR at this site. Lastly, we found that NAHR frequency varied depending on the primary orientation of cells in the individual, indicating a possible selection process influencing orientation frequency. Thus, by elucidating a method for quantifying NAHR with a new, greater level of precision, our results act as a pivotal first step to understanding, and ultimately treating the numerous diseases and disorders that are mechanistically linked to NAHR.

## Materials and Methods

Sperm samples were purchased from 2 sources, New England Cryogenics (n = 10), and California Cryobank (n = 10). Each sample set contained samples from Caucasian (n = 5), and African American or other ethnic group (n = 5) men. This was done in an effort to get an equal number of samples of each possible orientation as unpublished data from our group, as well as data from Giner-Delgado et al. (2019) suggests that a preponderance of individuals with a European heritage have the plus orientation while more of those of African and Asian descent have the minus orientation. DNA was isolated using the DNeasy Blood and Tissue Miniprep Kit (Qiagen). Cheek cell samples (n = 5) were collected to serve as controls, one from a known heterozygous female as a positive control for both assays, 2 from Caucasian men known to be of plus orientation, and 2 from men of African or Asian descent known to be of minus orientation. These cheek cells were collected using Epicentre Illumina Catch-All Sample Collection Swab Soft Pack and DNA extracted using Epicentre Illumina Masterpure DNA kit according to manufacturer protocol.

The digital PCR (dPCR) assay was done using the iPCR (inverted PCR) protocol found in Kirby et. al. (2016) but with shorter incubations times for Restriction digests and Ligation. In addition, a second restriction digest with SfiI was added to linearize the self-ligated loops. The custom primer/probe assay was provided by BioRad, but Thermo Fisher’s QuantStudio 3D Digital System was used for the Digital PCR according to recommended procedures. Details of entire protocol can be found in Supplemental Materials (S1 Appendix).

All California Cryogenic samples were run twice for technical replication, and resultant counts were averaged. There was not enough sample from New England Cryogenics for this control. In addition, a sampling of amplicons from each primary orientation was gel extracted and sequenced to verify fragment identity.

## Results

Any orientation count outside of a sample’s primary orientation is considered to be the result of NAHR. By taking the ratio of these NAHR counts over the total number of counts, we can calculate the frequency of NAHR. NAHR was detected in all samples, including the cheek cell controls, at a much higher rate than anticipated based on past literature. Averaged raw counts for each sample, averaged counts for each orientation per sample group, ratio of NAHR/Total calls for each sample, and the average of this ratio for each sample group, are presented in Table 1. Samples were grouped based on cell type (cheek versus sperm) and primary orientation (which orientation had the highest number of calls for that sample, plus or minus).

**Table 1.**
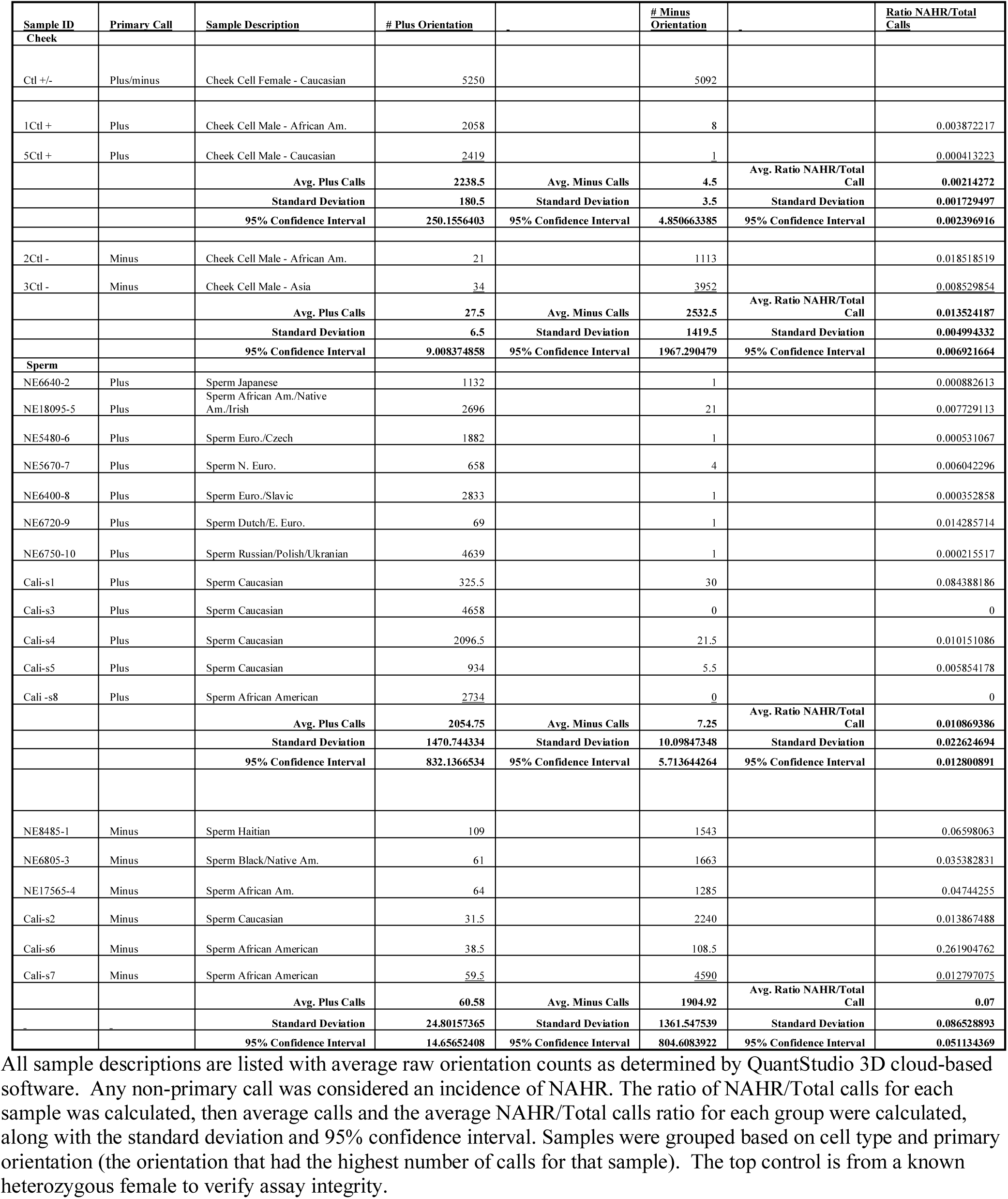
Samples, raw counts, and calculations of results.

Data from Table 1 are visualized in the following figures. Fig 2 displays the primary orientation calls for each sample, while Fig 3 displays the alternative orientation calls (considered NAHR) for each group. The overall number of primary calls is similar among all groups, ranging from 2058-2419 for Plus Cheek cells, 1113-3952 for Minus Cheek Cells, 69-4658 for Plus Sperm Cells, and 108-4590 for Minus Sperm Cells (Fig 2). However, alternative calls between groups are far more variable with a range of 1-8 for Plus Cheek Cells, 21-34 for Minus Cheek Cells, 0-30 for Plus Sperm Cells, and 31-109 for Minus Sperm Cells (Fig 3).

**Fig 2.**
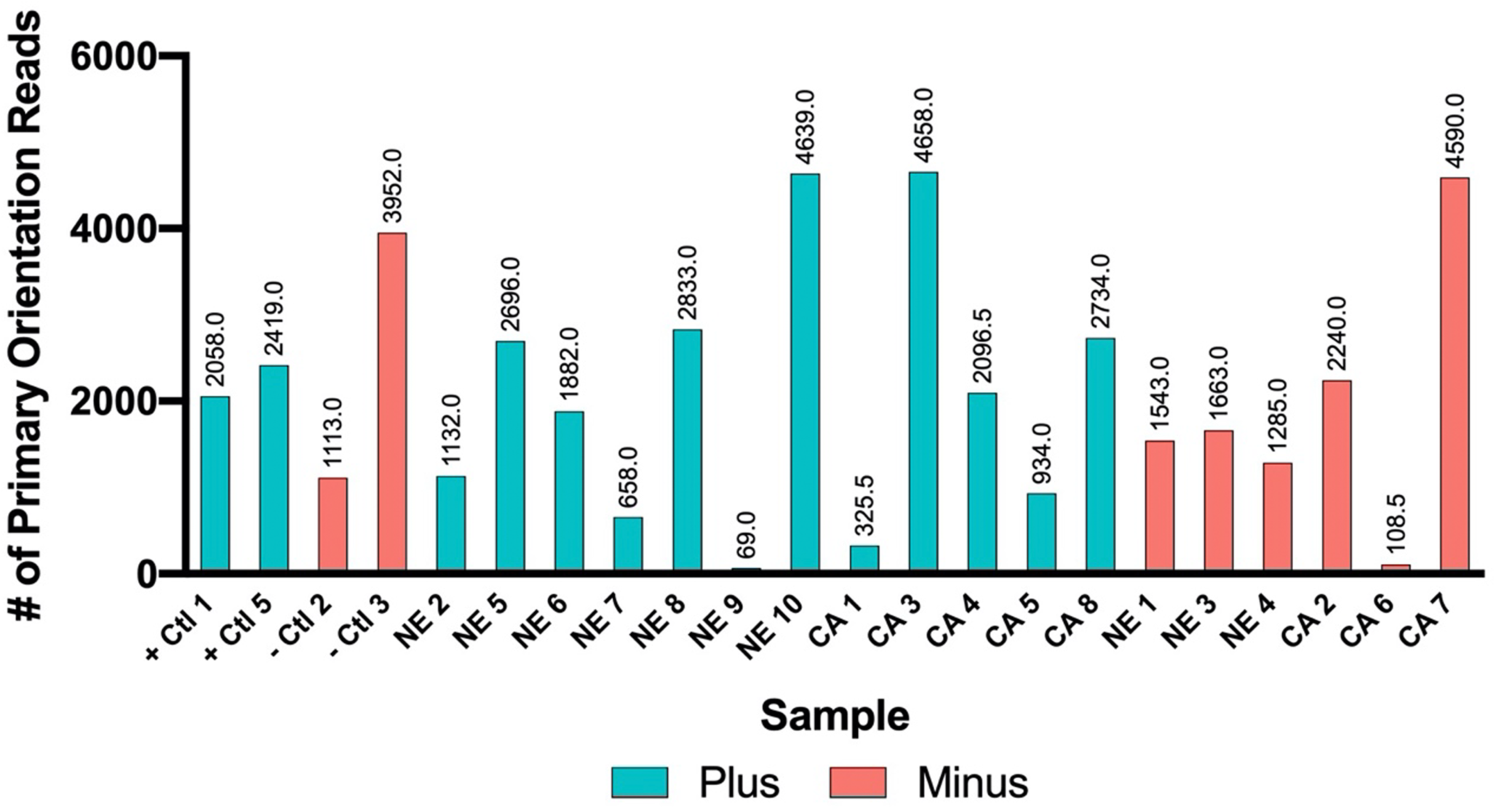
Bar graph of primary orientation counts for each sample, seen in Table 1. Orientation counts produced by digital PCR runs using QuantStudio 3D analysis. Plus calls are depicted in teal while Minus calls are in orange.

**Fig 3.**
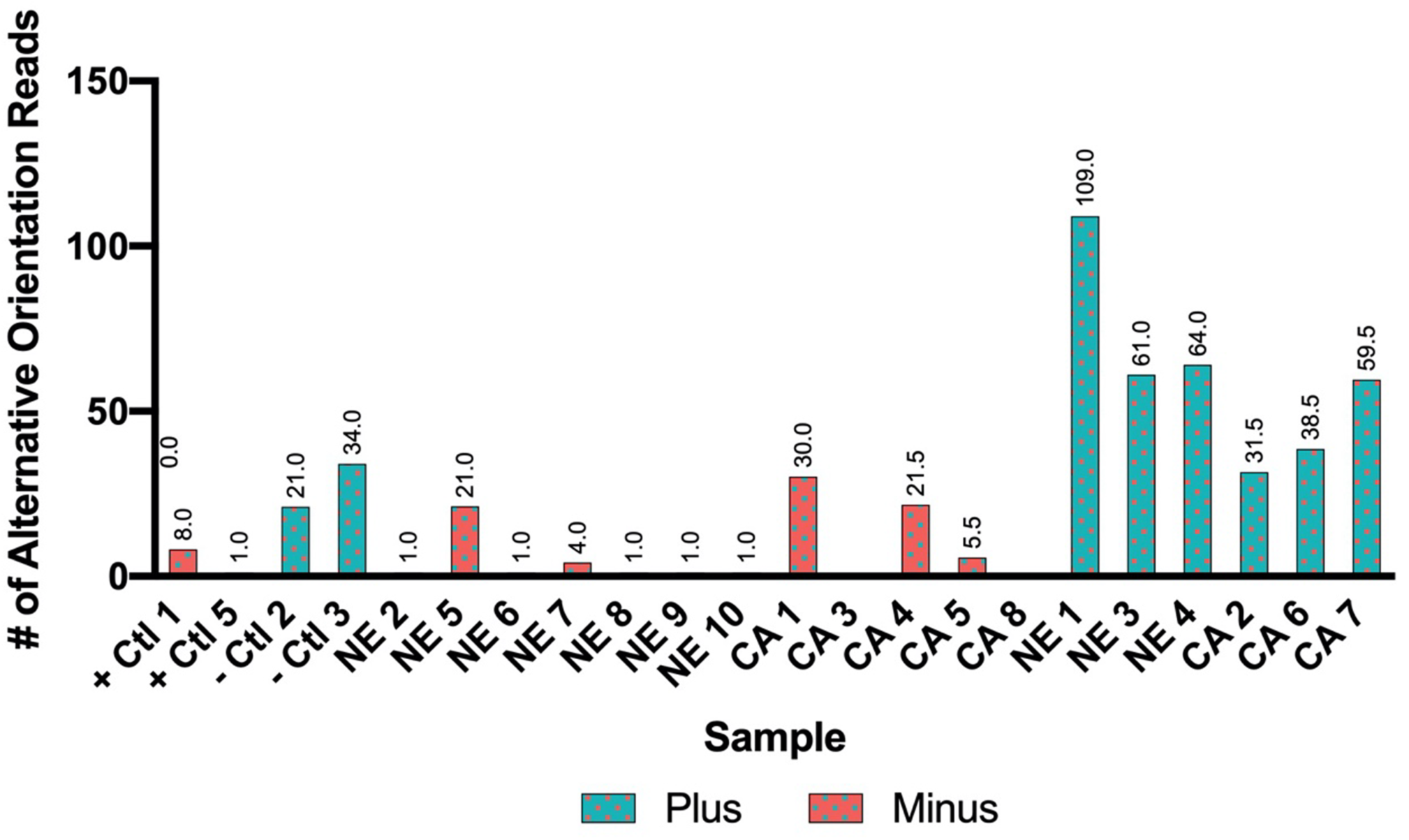
Bar graph of alternative orientation calls seen in Table 1. Calls produced by digital PCR runs using QuantStudio 3D analysis. Plus calls are in teal while Minus calls are in orange.

The NAHR frequency for these samples, calculated in Table 1 as the number of minor/alternative calls divided by the total number of calls, is visualized in Fig 4. NAHR frequency varies between groups ranging from .04-.39% for Plus Cheek Cells, 0.8-1.8% for Minus Cheek Cells, 0-8.4% for Plus Sperm Cells, and 1.3-26% for Minus Sperm Cells. Mean NAHR frequency can be seen in Table 1 for these samples with 0.2% in Plus Cheek cells, 1.3% in Minus Cheek cells, 1% in Plus Sperm cells, and 7% in Minus Sperm cells.

**Fig 4.**
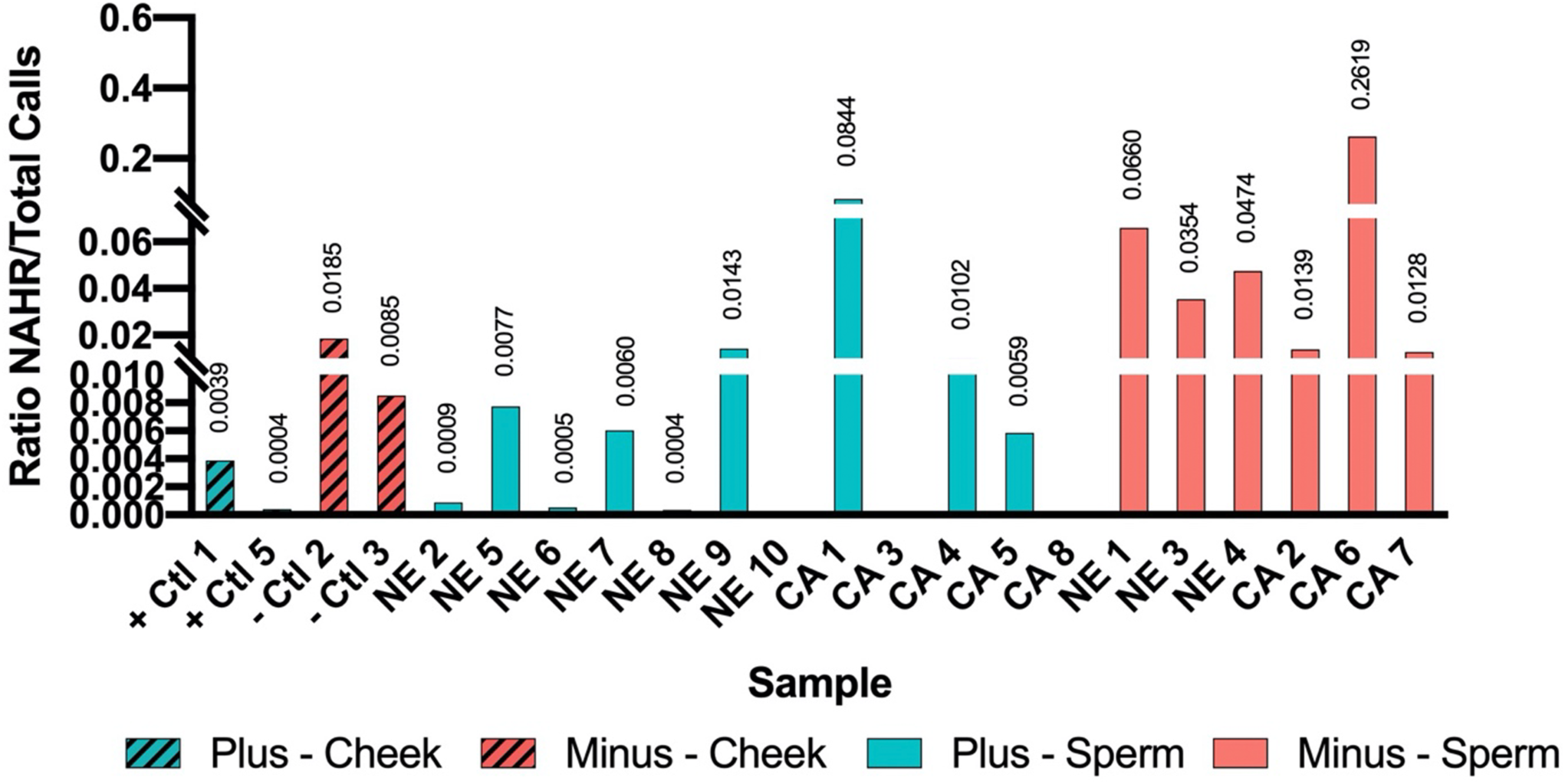
Calculated ratio of NAHR/Total calls for each sample. Calls produced by digital PCR runs using Quant-Studio 3D analysis. All non-primary orientation calls are considered indices of NAHR. Each primary-orientation/cell-type group is in its own color: Plus Cheek cells in hashed teal, Minus Cheek cells in hashed orange, Plus Sperm in teal, and Minus Sperm in orange. Note that CA-1 and CA-6 Sperm samples are outliers due to low total number of counts.

The difference in frequency of NAHR in each cell type based on primary orientation can be most easily viewed in Fig 5, where the ratio of NAHR/Total Calls is overlaid for each cell type and color coded based on primary orientation, or call. Here it can be seen that NAHR, in nearly all samples with a plus orientation, is lower than NAHR in all samples with a minus orientation, regardless of cell type.

**Fig 5.**
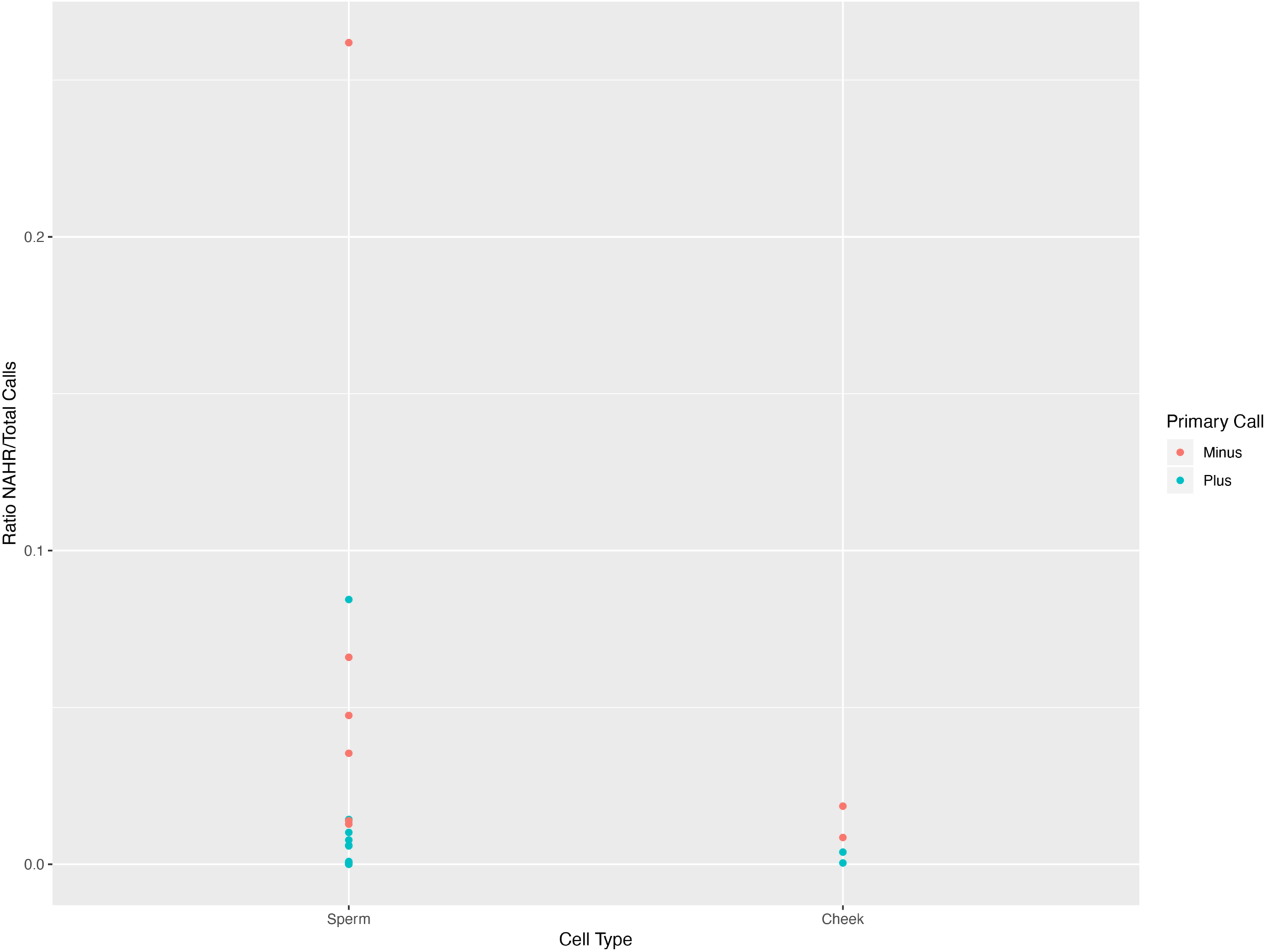
Plot of NAHR frequency based on cell type and primary orientation. Values for NAHR frequency for each sample overlaid for each cell type and color coded for primary orientation with Plus in teal and Minus in orange.

An ANOVA analysis and a multiple regression model fit to this data, verified that the difference in NAHR frequency based on primary orientation is significant (p = 0.04). In addition, a much larger variation in NAHR frequency can be seen for sperm versus cheek cells (Fig 5), with NAHR frequency in sperm cells at 1% for Plus cells and 7% for Minus cells, and in cheek cells at 0.2% for Plus cells and 1.3% for Minus cells. However, the broad range in the sperm is created by a few outliers that had low total calls, while the bulk of the sperm sample NAHR frequency lies within the same range as the 4 cheek cell samples. Thus, the difference in NAHR frequency based on cell type was not significant in the ANOVA (p = 0.41), and the multiple regression (p = 0.28), likely due to high standard deviation caused by technical differences in sample concentrations and some low sample numbers, particularly for cheek cells. Future study should include a larger number of cheek cell samples, treated as experimental samples versus controls in order to determine whether the NAHR seen in sperm cells reflects somatic mosaicism or the additive effects of mosaicism and meiotic NAHR.

## Discussion

Based on observations made during classroom genotyping of cheek cells for this Xq28 inversion, and Giner-Delgado et al. (2019), who identified 28 inversions, including this one, for which the minus orientation is more common in individuals of African descent, we sought to get reasonably balanced sample sizes for each orientation based on ancestral background of the samples. In addition, we utilized only males for each group to limit complexity and possible recombination events, looking at only a single X chromosome per individual, with the exception of the single heterozygous female used strictly to verify the validity of the assays to detect both orientations equally. The clear difference in magnitude of alternative calls (NAHR) for each group can be seen in Fig 3 compared with Fig 2, which shows a relatively similar trend of call numbers for primary calls in each group. Normalized NAHR calls as a ratio over total calls for each sample, depicted in Fig 4, shows clearly that NAHR is occurring in all sample types. Our observance of the inversion in cheek cells indicates that this type of NAHR with large inverted repeats is happening in somatic cells during mitosis, as proposed by Flores et al. (2007). However, our frequency differences between cell types was not significant, so it cannot be confirmed that NAHR occurs in sperm cells, as the NAHR observed there could be due to mosaicism alone.

Interestingly, NAHR frequency is significantly higher, according to our ANOVA analysis and multiple regression model, where minus is the primary orientation versus plus (Fig 5), indicating some sort of preference to invert to the plus orientation. This may be a regional effect. In 2005, Steffanson et al. noted a similar effect at the 17q21.31 region with the discovery of a 900kb inversion, elucidating a key source of variation in the human genome and its organization, contributing to the divergence of the H1 and H2 haplotype lineages. Their study indicated that one orientation was associated with the H2 lineage, more common in European populations, but rare in Africa and Asia, versus the H1 lineage which houses the other orientation for this inversion. Because it is so rare in Africa, the H2 lineage is assumed to have grown from a few founder chromosomes in groups that left Africa over 60,000 years ago, and due to some positive selection on the H2 haplotype structure in the local environment of Europe, became the prominent haplotype. Kirkpatrick et al. (2006) supported this idea of inversions utilized in local adaptation, pointing to adaptive, intraspecific variation involving inversions in Drosophila, Rhagoletis, and Coelopidae flies, as well as Anopheles mosquitoes. Flores et al. (2007) provided further evidence in finding a higher number of inverted structures in adult, versus newborn, blood cells for all three inversions in their study. Thus, the local adaptation mechanism may be at play here with this inversion as well.

It has been reported that 40 X-linked diseases have been mapped to the Xq28 region (Kolb-Kokocinski et al. 2006). Thus, understanding how often this inversion occurs, and the possible affects it has on surrounding gene expression could provide valuable insight into some of these conditions. Giner-Delgado et al. (2019) in a large-scale data mining of various databases, determined that the HsInv0389 inversion is linked to expression differences of several genes in various tissues, all of which are located in the Xq28 region. Based on these findings, they further speculated that the recurrence of this inversion in mammalian evolution, observed by Caceres et al. (2007), may reflect selective pressure on the two different orientations.

Using digital iPCR, this study determined a way to accurately genotype the HsInv0389 inversion, located at the end of the q arm of the X chromosome in individual males with different primary orientations. Our findings indicate that this technique is a robust and accurate approach to quantitate the frequency of NAHR at this site, and is readily modifiable for similar sites. Digital iPCR verified that NAHR is happening at a much higher frequency than had been previously determined, and the assay detected inversion generation, within the individuals, in both germ-line and somatic cells. The results indicate a favoring of NAHR from Minus to Plus, versus Plus to Minus for the samples in this study that were all collected in North America. Inversion specific haplotypes develop as a result of suppressed recombination and Steffanson et al. (2005) report on two inversion haplotypes on chromosome 17 that show no evidence of recombination over a possible 3 million years of divergence. NAHR at the frequencies detected in this study should lead to the breakdown of such strong linkage disequilibrium through the movement of haplotypes from one gene arrangement to the other and subsequent recombination in homokaryotypes. In addition, it would be interesting to test the local adaptation theory of Steffanson et al. (2005) and Kirkpatrick et al. (2006) by conducting this study in parts of the world where the primary orientation for the majority of the population is Minus to see if the direction of NAHR frequency varies from this study.

## Notes

### Competing Interest Statement

The authors have declared no competing interest.

