## Supplementary material for "Characterizing the frequency of non-allelic homologous recombination (NAHR) on the human X chromosome (Xq28) using digital inverted PCR": S1 Appendix

**S1 Appendix. Digital iPCR Protocol**

**Restriction Enzyme Digest Reaction**

1. A master mix was made for all samples whereby each sample received:
   - 20 µL BglII Buffer (NEB cat # R0144S)
   - 179 µL $H_2O$ and DNA (1µg of DNA as determined by nanospec)
   - 1 µL BglII Enzyme (NEB cat # R0144S)

2. Digestion reactions were built in 0.5ml tubes and placed in a PCR machine programmed as follows:
   - 6 hours at 37$^{o}$C (1µL BglII spiked in at the 3-hour mark)
   - 20 minutes at 85$^{o}$C
   - Hold at 4$^{o}$C until collected

**Self-Ligation**

1. A master mix was made for all samples whereby each sample received:
   - 60 µL T4 DNA Ligase Buffer (NEB cat #B0202S)
   - 338 µL $H_2O$
   - 2 µL T4 DNA Ligase Enzyme (NEB cat #M0202S)

2. Aliquots of 400µL of master mix were dispensed into 1.7ml tubes and the entire 200µL digest of each sample was added to its respective ligation tube and incubated at 25$^{o}$C for 2 hours, followed by 20 minutes of heat inactivation at 65$^{o}$C and then placed at 4$^{o}$C.

**Re-linearization with SfiI Digestion** to increase efficiency of the dPCR assay

Samples were changed back into water solute by running through Microcon 100 columns by Millipore (Cat#Z648094)

1. A master mix was made for all samples whereby each sample received:
    - 5 µL CutSmart Buffer (NEB cat # B7204S)
    - 40 µL $H_2O$ and DNA (all of DNA resulting from the Microcon column)
    - 5 µL SfiI Enzyme (NEB cat #R0123S)

2. Tubes were incubated for 1 hour at 50ºC
3. Samples were then put through the same Microcon 100 column to re-purify back into water to insure purity for the dPCR reaction.
4. Samples were quantitated on a Nanodrop 8000.

**Digital PCR**

Primer probe assays were custom created by BioRad as follows – note that the reverse primer for the Plus reaction is the forward primer for the Minus reaction:

Plus Reaction:

Product: PrimePCR Custom Assay Concentration: 20x

Control: 195549772

Item: 10031276

Format: 200 x 20 µL reactions

Assay Info: dPCR Plus

Forward: CACATAAGCCACACCACAGG Reverse: TCATCACGCGAAAAACAGAG Probe: AGCCGTCCTTGCGGTCCTC Primer:Probe: 900 nM:250nM

Dye Quencher: 5' 6-FAM, 3' Iowa Black® FQ

Minus Reaction:

Product: PrimePCR Custom Assay Concentration: 20x

Control: 202473776

Item: 10031279

Format: 200 x 20µL reactions

Assay Info: dPCR Minus

Forward: TCATCACGCGAAAAACAGAG Reverse: TGCATCCCCTCTAGTCGAA Probe: CCGCCGCCTCTGGGTCTC Primer:Probe: 900 nM:250nM

Dye Quencher: 5' HEX, 3' Iowa Black® FQ

1. Reactions were built by adding 90ng of DNA, 0.75µL of each of the above assay mixes, 7.5µL of a 2X QuantStudio 3D Digital PCR Master Mix buffer provided by ThermoFisher cat#A26358, and brought up to 15µL with water. Each sample was loaded onto a 3D Digital 20K PCR chip (ThermoFisher cat#A26316), and run on a ProFlex PCR system with the following program:

- 98°C 5 minutes
- 39 cycles
    - 98°C 1 minute
    - 53°C 30 seconds
    - 60°C 90 seconds
- 60°C 2 minutes
- 10°C Indefinitely

Chips were then visualized on a QuantStudio 3D Chip reader and probes were counted for each reaction with cloud-based QuantStudio 3D software.
